# Metabolic activity organizes olfactory representations

**DOI:** 10.1101/2022.07.21.500995

**Authors:** Wesley W. Qian, Jennifer N. Wei, Benjamin Sanchez-Lengeling, Brian K. Lee, Yunan Luo, Marnix Vlot, Koen Dechering, Jian Peng, Richard C. Gerkin, Alexander B. Wiltschko

**Affiliations:** Google Research, Brain Team; Cambridge, MA, USA; Department of Computer Science, University of Illinois; Urbana-Champaign, IL, USA; TropIQ Health Sciences, Netherlands

## Abstract

Hearing and vision sensory systems are tuned to the natural statistics of acoustic and electromagnetic energy on earth, and are evolved to be sensitive in ethologically relevant ranges. But what are the natural statistics of *odors*, and how do olfactory systems exploit them? Dissecting an accurate machine learning model^1^ for human odor perception, we find a computable representation for odor at the molecular level that can predict the odor-evoked receptor, neural, and behavioral responses of nearly all terrestrial organisms studied in olfactory neuroscience. Using this olfactory representation (<u>P</u>rincipal <u>O</u>dor <u>M</u>ap, POM), we find that odorous compounds with similar POM representations are more likely to co-occur within a substance and be metabolically closely related; metabolic reaction sequences^2^ also follow smooth paths in POM despite large jumps in molecular structure. Just as the brain’s visual representations have evolved around the natural statistics of light and shapes, the natural statistics of metabolism appear to shape the brain’s representation of the olfactory world.

## Intro

Sensory neuroscience depends on quantitative maps of the sensory world. Color mixing principles^3–5^ and corresponding biological mechanisms^6–8^ help explain the organization of color perception in the early visual system. Gabor filters describe the receptive fields of visual cortex (V1) simple cells in later stages of visual processing^9^. They also account for the organization of acoustic energy from high-to low-frequency and explain the tonotopic representation of perceptual tuning in animal hearing^10,11^. Understanding these sensory representations is critical for the design and interpretation of experiments that probe the organization of our sensory world. However, a representation and organizational framework for odor has not yet been established. Even though structure-activity relationships in human olfaction have been explored^12–15^, “activity cliffs” – seemingly small changes in molecular structure that produce profound changes in activity^16^ (such as odor) – have limited the generalizability of representations developed from structural motifs^14^. Does such a representation for odor – common to species separated by evolutionary time – even exist?

Machine learning models, particularly neural networks, have identified common representations encoded in biological nervous systems for sensory modalities, including vision and audition^17^. For instance, the first few layers of convolutional neural networks – trained on visual scenes drawn from natural statistics – learn to implement Gabor filters^18^. More strikingly, learned representations at progressively deeper layers of neural networks predict the responses of neurons in progressively deeper structures in the ventral visual stream^19,20^. Similarly, neural networks trained to classify odors can also match olfactory system connectivity^21^. Representations of the sensory world learned by training predictive models thus often recapitulate nature.

Here we perform a comprehensive meta analysis on 12 olfactory neuroscience datasets^12,22–31^, spanning multiple species and levels of neural processing. We find that the embedding from a graph neural network trained on human olfactory perception, which we term the *principal odor map* (*POM*), is highly predictive of the olfactory responses in nearly all datasets, even for species separated by hundreds of millions of years in evolution. In addition, we find POM is specific to olfaction as it shows no advantage in enteric chemoreception tasks or the prediction of general physico-chemical properties. The existence and specificity of POM not only suggest a shared representation of odor across animals but also provide an accurate and computable framework to study the organization of odor space. We show that metabolic reactions that determine the states of all living things – and the odors they emit – explain the organization of POM, and that multi-step reaction paths are smooth trajectories in POM. Finally, we identify strong associations between POM and the natural co-occurrence of molecules in natural substances. Together, these results suggest that the natural statistics of biologically-produced molecules shaped the convergent evolution of animal olfactory systems and representations despite significant differences in biological implementation.

## Results

### Neural network embedding as a principal map for animal olfaction

Graph neural networks (GNNs) show state-of-the-art ability to accurately predict human olfactory perceptual labels in both retrospective^41^ and prospective settings^1^. Here, we use a GNN embedding – the neural network layer immediately preceding the task-specific architecture – as a representation of odor and evaluate its predictive power in a meta analysis across 12 datasets (Methods) in olfactory neuroscience spanning 9 common model species, including mosquito, fruit fly, and mouse, as well as different scales of biology, including olfactory receptor, neuron response, and whole-animal behavior (Figure 1a; Extended Data Figure 1). We quantify the predictive performances of the GNN embedding on regression or classification tasks for these curated datasets, and compare its performance against generic chemical representations often used in the predictive chemoinformatic models. As shown in Figure 1b, the embedding predicts receptor, neural, and behavioral olfaction data better than generic chemical representations across species separated by up to 500M years of evolution – and possessing independently evolved olfactory systems. We thus term this embedding the principal odor map, or POM.

**Figure 1:**
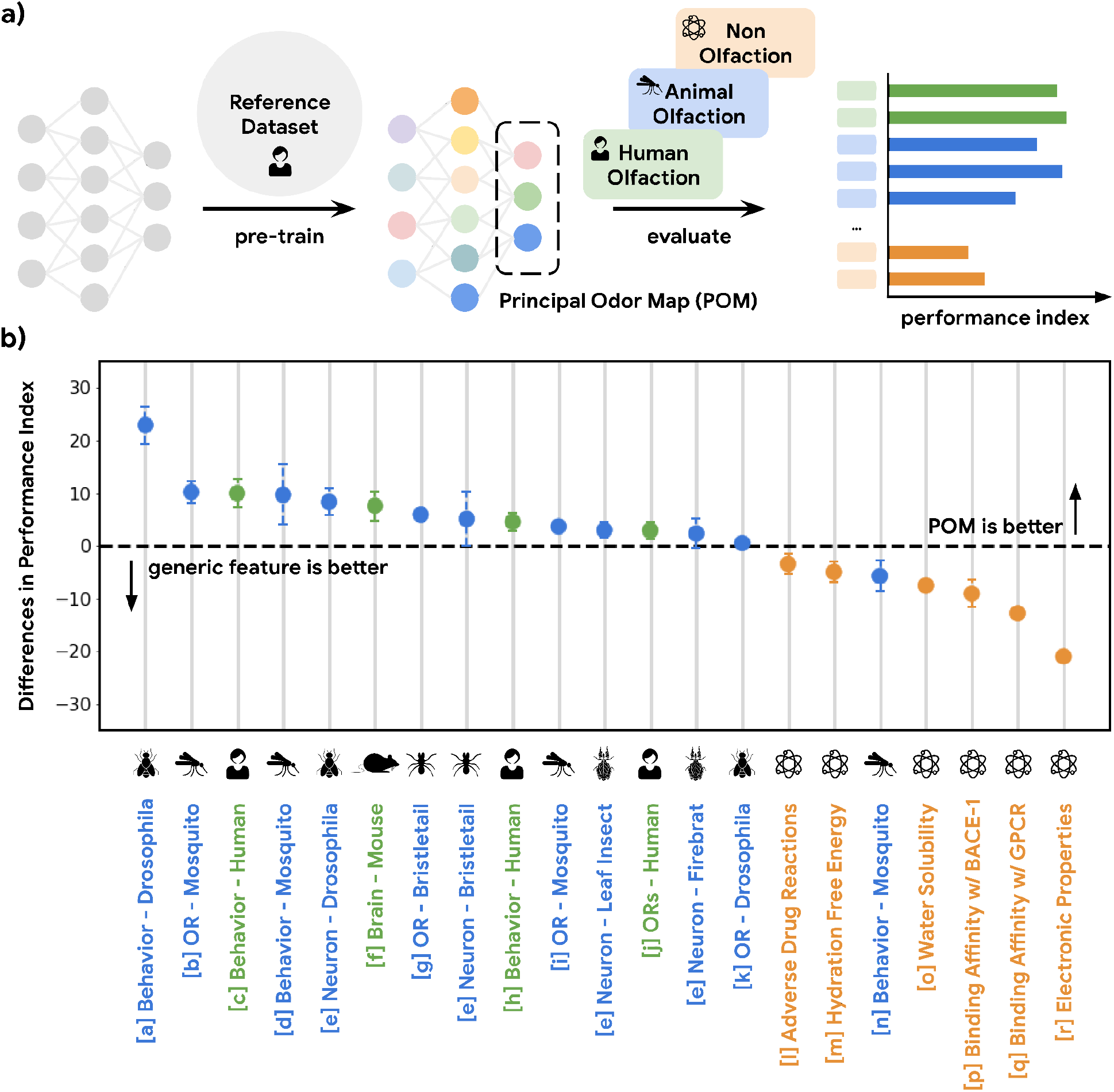
A single latent space can explain olfactory data across species and scales. **a)** A graph neural network model pre-trained on human olfactory perceptual data produces a principal odor map, or POM (latent space, dashed box), which can be used to make predictions about any small, volatile molecule in biological and behavioral experiments. **b)** A random forest model using only POM produces predictions that meet or exceed those obtained from commonly-used generic molecular features^32,33^ (Mordred) across a range of olfactory datasets^12,22–31,34–40^ in different species (green for vertebrates and blue for invertebrates), but not for prediction of non-odorous molecular properties (orange). The Y-axis is the difference between performance indices for models using POM vs. generic molecular features. Performance index is a rescaled metric to place classification and regression performance on the same axis. Performance indices of 0 and 100 represent random and perfect predictions, respectively. Error bars are calculated as the standard deviation of performance differences across multiple random seeds.

### The principal odor map is specific for olfaction

While the POM exhibits generalizability across olfactory tasks in various species, it should be no better than generic chemical representations on tasks irrelevant to olfaction in order to optimize its representational power specifically for olfaction (i.e. the no-free-lunch theorems^42^). As shown in Figure 1b, POM does not show a significant or consistent advantage over generic chemical representations for predicting molecular properties that are not likely exploited by olfaction, such as electronic properties (e.g., QM7^39^) and adverse drug reactions (e.g., SIDER^35^) compiled by MoleculeNet^34^. We then apply POM to predict molecular binding activity for G-protein-coupled receptors (GPCRs, of which mammalian olfactory receptors are only a subset^29^) generally, including those involved in enteric chemical sensation^40^ (e.g., 5HT1A for serotonin and DRD2 for dopamine). While POM demonstrates superior performance for GPCRs involved in human olfaction, their performance is significantly worse for GPCRs related to enteric chemical sensation compared to generic chemical representations, showing specificity for olfaction (Figure 1b, Extended Data Figure 2). We observe a similar result when we restrict the analysis to only the original training molecules, showing that it is the task and not the molecule which determines the suitability of the POM. (Extended Data Figure 3).

### Metabolic activity explains the organization of the principal odor map

Since animals have different biological implementations for external molecular detection (e.g., ionotropic receptors for mosquitoes and independently evolved metabotropic GPCRs for mammals), it is surprising that a human-derived representation of odor can explain responses in a diverse set of species. We hypothesize that such convergent evolution could be the result of a shared natural environment for most animals where they experience the same set of ethological signals, including various nutrients and pheromonal cues from metabolic processes; in other words, detecting and identifying the state of living things by their odor is broadly useful across species.

To test this hypothesis, we explored all odorant molecules in a carefully curated metabolic reaction database called MetaCyc^2^, containing experimentally elucidated reaction pathways. We identified 17 species with sufficient metabolic data, spanning 4 kingdoms of life (Extended Data Figure 4). We then constructed networks of metabolites for these species in which directed edges represent the direction of a reaction between one node (a reactant) and another (a product) (Figure 2a). We then computed the discrete “metabolic distance” between any two compounds by calculating the shortest paths through these networks (Figure 2b; Extended Data Figure 4). From those metabolic networks with enough metabolites (>100), we repeatedly sampled 50 pairs of odorants (molecules that pass a validated rule set for odor probability^43^) for each metabolic distance ranging from 1 to 12, and asked how well the distance in POM correlates with these metabolic distances (Figure 2c). We found that there is a strong correlation between the metabolic distance and POM distance (r=0.93), and that common measures of structural similarity between these metabolites can only account for part of the relationship (r<0.8). This is especially true for neighbor metabolites in metabolic networks where a biological reaction changes the molecular structure drastically; while such drastic changes produce large structural distances, only smaller changes in POM are observed -- the reactant and product spanning this structural cliff frequently share a common odor profile (Figure 2d).

**Figure 2:**
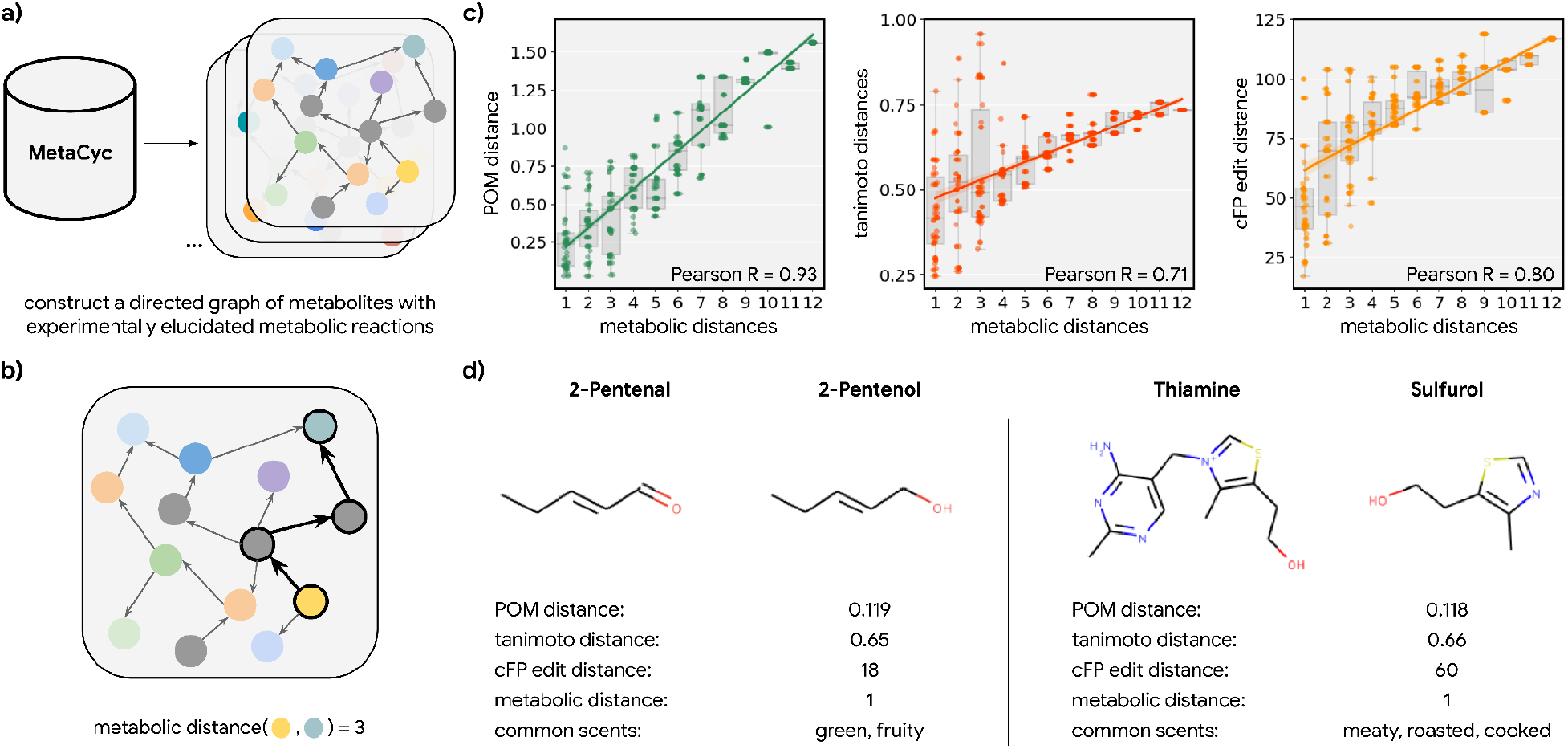
Metabolic pathways predict distance in the principal odor map (POM). **a)** The contents of MetaCyc, a large database of experimentally-elucidated metabolic reactions across multiple species, were used to construct directed graphs connecting metabolites, including those with odors (non-gray). **b)** The discrete pairwise distance of two molecules was defined by the shortest directed path between them within a species’ metabolic graph (if any). Each step corresponds to a single chemical reaction specified in MetaCyc. **c)** Continuous pairwise distances between molecules in POM – which was produced from human perceptual data alone – are strongly correlated with discrete metabolic distance (left, r=0.93). This effect is not driven solely by the structure similarity of related metabolites, since a weaker relationship is observed using alternative structural distance metrics including tanimoto distance (center, r=0.71) and edit distance in count-based fingerprints (right, r=0.80). **d)** Two pairs of example molecules that are closely related in metabolism. While these are structurally dissimilar molecules (tanimoto distance>0.65; left: change in a key functional group; right: removal of a major substructure), a single metabolic reaction can turn one to the other, and therefore, POM also organizes them closely together (POM distance<0.12). In turn, they have similar odor profiles.

Having established that metabolic distance was closely associated with distance in POM, we next asked whether metabolic reactions are easier to understand and interpolate in POM. If a pathway of reactions proceeds in a consistent direction in a molecular representation, then that pathway can be identified with that direction (e.g., “toward fermentation”); alternatively, the pathway could simply be a random walk in space. Using principal components analysis, we visualized the metabolic pathway for both DIBOA-glucoside biosynthesis (Figure 3a) and gibberellin biosynthesis (Figure 3b) in 2D with both count-based structure fingerprints (cFP) and POM. We find the organizations of POM show a smooth progression from starting metabolites to final product metabolites, even though the same pathways show irregular progressions when organized simply by molecular structure. To further quantify such effects, we examine the “smoothness” of all triplets of three consecutive metabolites (pairs of consecutive reactions) in 37 unique metabolic pathways with only odorant molecules (Figure 3c). As shown in Figure 3d, most of the paths for these triplets become smoother after the pre-trained neural network projects their structure into POM (see Extended Data Figures 7 and 8 for additional controls and analysis), suggesting that the organization of POM reflects a deeper relationship between olfaction and metabolic processes.

**Figure 3:**
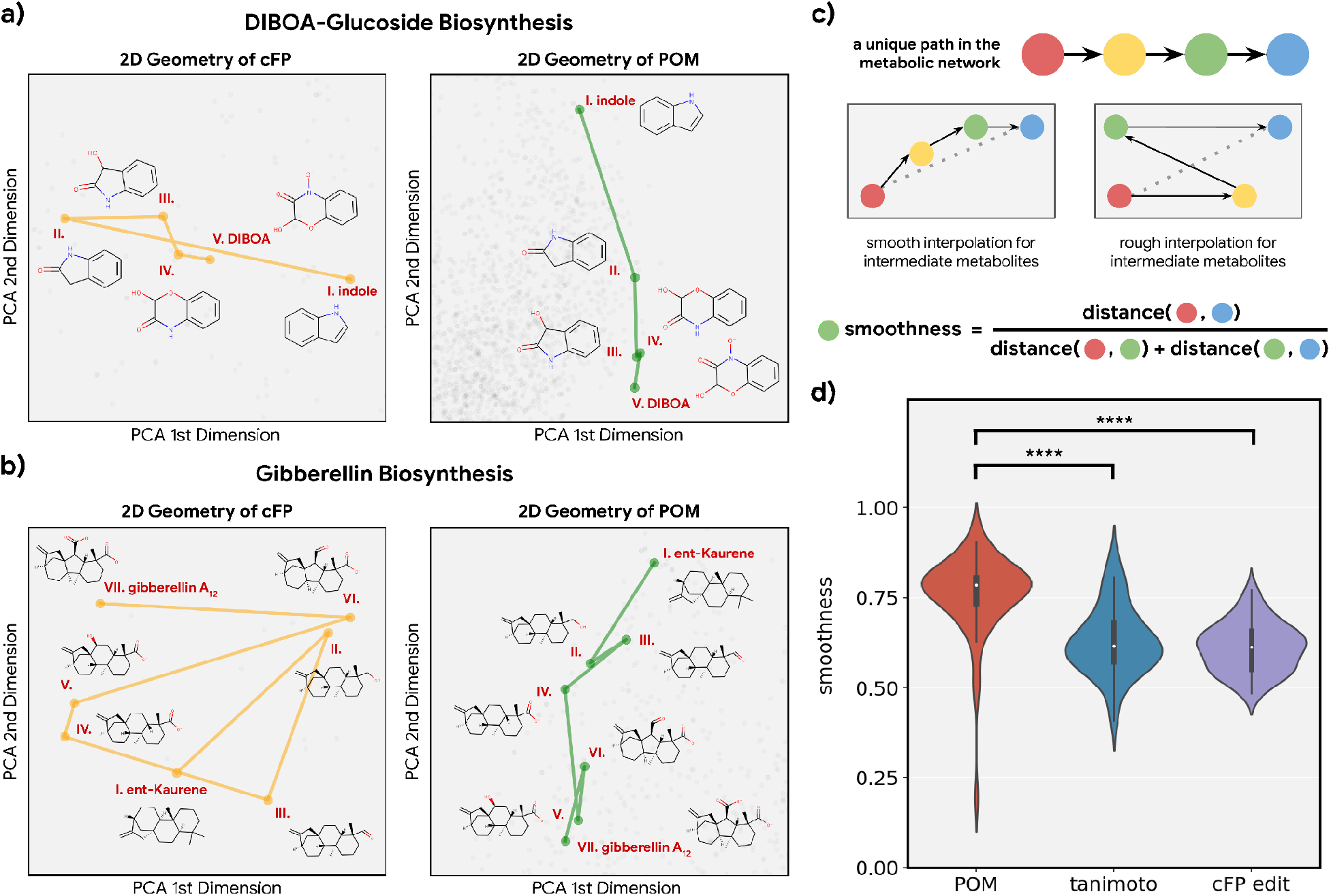
Smoothness of metabolic pathways in the principal odor map (POM). **a)** Left: A 4-step pathway (DIBOA-glucoside biosynthesis) depicted in a 2D representation of the structure fingerprints (count-based fingerprints, or cFP) using principal components analysis. Right: the same pathway depicted in a 2D representation of the principal odor map (POM). **b)** Left, the same 2D cFP representation for a 6-step pathway (gibberellin biosynthesis). Right, a 2D representation of the same pathway in the POM. We observe relatively smooth trajectories in POM for these pathways even though the same pathways show irregular trajectories in the structure space. **c)** To systematically quantify such “smoothness”, we examine all unique pathways in the metabolic network (top). A desirable molecular representation should exhibit smooth reaction paths, proceeding in a more consistent direction from the starting metabolite to the final metabolite allowing interpolation for intermediate metabolites (center). Smoothness for an intermediate metabolite is formally defined as the ratio between the direct euclidean distance and total path length between the start and end metabolites. A smoother path will result in a ratio close to 1 (bottom). **d)** Metabolic trajectories are smoother after metabolite structures are projected to POM than when using alternative structural distance metrics (paired t-test, p<0.0001 for both structure distance metrics).

### Molecules that co-occur in nature are also closer in the principal odor map

To further validate our hypothesis, we investigated 214 molecules that co-occur in 303 essential oils aggregated in the Pyrfume data repository^45^. Molecules which co-occur in the same object in nature usually convey similar ethological cues, including danger, conspecifícs, or in the case of plants, nutrient availability. If the organization of POM is indeed driven by the shared natural ethological signals driven by metabolic processes, they should also be represented similarly in the POM.

We calculated the distance for all molecule pairs in POM and also using a common structural similarity metric (Figure 4a). If co-occurring pairs share metabolic origins, then such co-occurring pairs should be closer together in POM than would be expected by chance. Indeed, the distance distribution for co-occurring molecule pairs is shifted towards “nearness” in POM; this shift is larger than one would expect when only considering their structure similarity (Figure 4b,c). As an illustration, we sample two pairs of co-occurring molecules – one pair from rose oil and the other from orange oil – that have a very distinct within-pair structure (tanimoto distance > 0.95, Figure 4d). Despite their distinct structures, we discover that both pairs of molecules are part of the terpenoid biosynthesis pathway and are closely related downstream products of geranyl diphosphate (Figure 4e). The POM representation – in which the distance within each such pair is small – captures their physical co-occurrence and proximity in the metabolome.

**Figure 4:**
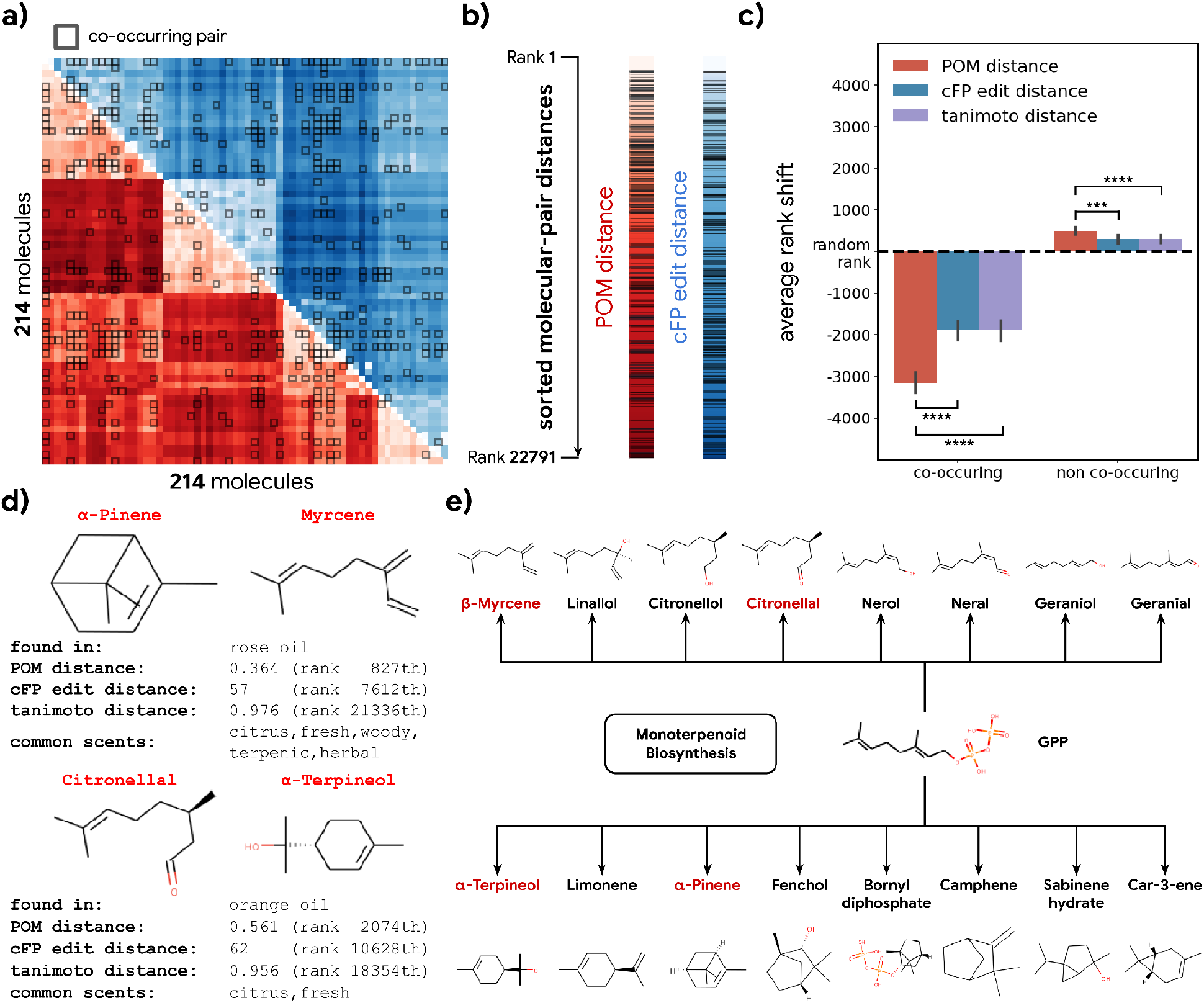
Co-occurrence of odorous molecules in natural substances is explained by the principal odor map (POM). **a)** We compiled a dataset of 214 molecules found in 303 essential oils and computed their pairwise POM distance (red) and cFP edit distance (blue), where molecule pairs that co-occur in the same essential oil are indicated by dark boxes. **b)** To make POM and cFP edit distance comparable, we sort both POM and cFP edit distances for all 22,791 molecular pairs in the dataset from small to large, and mark co-occurring pairs with dark lines; we then **c)** plot the average shift in distance rank (relative to a random pair) for co-occurring (left) and non co-occurring (right) molecule pairs under POM distance (red), cFP edit distance (blue), and Tanimoto distance (purple). As expected, the co-occurring pairs have a smaller rank (nearer together) and non co-occurring pairs have a higher rank (further apart). More importantly, this rank shift for co-occurring molecules is ~2x larger in POM than for structures distance (paired t-test, p<0.0001), and reversed for non-co-occurring pairs (paired t-test, p<0.001). Error bar indicates the 95% confidence interval. **d)** Two example pairs of co-occurring molecules that POM successfully recognizes as closely related while conventional structure-based distance fails. Common odor labels for the two molecules as predicted by a state-of-the-art model^41^. **e)** The terpenoid biosynthesis pathway shows that these molecule pairs (red) are close downstream metabolic products of geranyl diphosphate^44^, explaining both their co-occurrence and their proximity in the POM despite their dissimilar structures.

## Discussion

In this study, we found that the embedding of a graph neural network pre-trained on a reference human olfaction dataset can be used as a principal odor map (POM) for predicting general receptor, neuron, or animal behavioral olfactory tasks across a large number of datasets. Using POM as an accurate and computable proxy for odor representation, we then proposed a hypothesis for the organization of odor with three facets: (1) Co-occurring molecules in nature are also nearby in the POM; (2) the underlying metabolic network dictating co-occurrence is highly correlated with the POM representation; and (3) directed metabolic reaction pathways trace smooth and consistent paths in the same representation. These results suggest that evolution of numerous terrestrial species’ olfactory systems has converged to decode a shared and principal set of ethological signals organized by the metabolic processes of nature and the natural statistics that emerge from it. Therefore, we hypothesize that the olfactory *umwelt* may be more similar across species than previously appreciated.

This claim unlocks new directions for olfactory neuroscience, animal ethology, and metabolomics. First, we predict that olfactory neural representations in most animals should be well-explained by POM or future representations based on similar principles. Theory suggests that tuning of neurons^46,47^ and even plasticity^48^ should reflect the natural statistics of odor; these statistics may be organized by metabolic activity, and may have esoteric geometries^49^. Second, the homology between odor space and metabolic space suggests that animal olfactory behavior may be broadly organized around the detection and discrimination of metabolic markers and even holistic metabolic states^50^: How ripe is this fruit^51^? How healthy is this mate? How nutritious is this soil? Third, the discovery of novel metabolic reactions and pathways could be informed by results from olfaction itself; molecules which have a similar smell – for reasons otherwise unknown – may be neighbors in an ethologically-relevant metabolic network. Mechanistically, olfaction mediated by the trace amine-associated receptors (TAARs)^52,53^, which detect metabolically-downstream products of essential nutrient amino acids, already shows the signature of a “metabolism detector”. Our results suggest this may be a feature of olfactory chemosensation more broadly. Over both evolutionary time and individual learning and development, the structure of metabolomes could also provide a substrate enabling smooth, separable manifolds to develop in neural activity space, a likely requirement for invariant object recognition^54^.

How universal can the principal odor map be, given that outputs of sensory neurons and downstream behaviors are driven by both evolution and learning? Certainly individual experiences can modify odor perception and act as an additional organizing force on any olfactory map. But environmental correlations are also learned through experience, and many of these are broadly shared across both individuals and species. For example, direct experience of the co-occurrence of citronellal and alpha-terpineol in an orange peel should reinforce, through olfactory plasticity in the brain (e.g., in cortex), those evolutionary changes in the periphery (e.g., receptors) driven by that same co-occurrence. Indeed, we found no clear pattern in relative predictive performance of the POM as a function of processing stage (from periphery to behavior). However, we do observe that the performance of POM improves non-asymptotically as a function of training data size and quality^41^; it is thus unclear where the ceiling lies for this approach.

In some networks, connected nodes are not only close in graph distance (e.g. one hop away) but also in physical distance (e.g. same household, same postal code, etc.) Similarly, we show that metabolic distance is closely related to odor distance. But there are surely exceptions; a single edge of a social network graph may span continents, and this exception to the general pattern may be important for explaining the macrostructure of the phenomenon. By analogy, such exceptions for odor, where a single metabolic step gives rise to a radically unrelated odor profile, could represent the boundaries between large, innate odor categories. Given the high correlation between metabolic distance and POM distance that we observe, these are indeed exceptions to a general rule. Future work, relying upon larger metabolic pathway datasets (especially including pathways that have yet to be elucidated), might find enough such exceptions to determine their meaning.

Ludwig Boltzmann and Erwin Schrödinger argued that the fundamental object of struggle for organisms is to identify and feed upon negative entropy (free energy)^55,56^. Each life form appears to be equipped with enzymatic and mechanical tools to access niches of free energy. Eleanor Gibson theorized that perception becomes refined during development to intuitively access this information^57^. Indeed, the sense of smell seems designed to identify quantities and accessibility classes of chemical free energy, which are determined in turn by metabolic processes in living things. Odors thus tend to be more similar within specific pockets of the chemical ecosystem and the carbon cycle. Evolution may have thus ethologically-tuned neural representations of chemical stimuli to the statistics and dynamics of this cycle.

## Methods

### Principal odor map

The principal odor map, or POM, corresponds to the activations of an embedding layer within a neural network, pre-trained on human olfactory perceptual data^41^. Briefly, the neural network contains three components: 1) a graph neural network (GNN) that represents the molecule as a graph and learns a representation for each atom through message-passing, 2) a multi-layer perceptron that aggregates the atom representations and learns an embedding for the entire molecule, and finally 3) a single fully connected layer predicting different odor descriptors. After pre-training the neural network, the parameters of the models are fixed, and the first and second components are used deterministically to generate the location of arbitrary odor-like molecules within the POM; equivalently, the molecule is represented as a graph and projected to a single vector representing the activations of the neural network’s final embedding layer. The representation of any odor-like molecule in the POM is this vector.

### Performance index for supervised learning

Under the supervised learning setting (using molecular featurization to predict out-of-domain results, see Figure 1), the performance index of a dataset for a specific representation (e.g., POM or Mordred^32^) is calculated from a random forest model’s performance using that specific representation as input features. The supervised learning setting includes some datasets with category labels (classification) and some with real number labels (regression).

For datasets of size N <= 200, we split the data in a leave-one-out fashion where we hold out one molecule at a time for evaluation and train with N-1 data points until all molecules have been held out once. For each split, N different seeds are used to initiate the model and the training data is jackknife resampled. For larger datasets of size N > 200, we perform a five-fold cross validation split of the data instead, and for each split, 100 seeds are used to initiate the model and the training data is resampled with replacement.

For datasets with categorical labels we compute auROC, while R^2^ score is compute for datasets with real number labels. To calculate the performance index, we then rescale the auROC or R^2^ score for each dataset such that 0 represents random performance and 100 represents perfect performance. Specifically, auROC can be converted to the performance index with (auROC - 0.5) * 100, and R^2^ score can be converted to the performance index by multiplying by 100. Finally, performance indices are averaged across all the seeds and targets (in those cases where the dataset contains multiple targets).

Since the optimal hyper-parameters for the model can be different for different representations and datasets, we perform a scan for important hyper-parameters for random forest models including number of trees in the forest, different ways to assign weight for each class label, number of features to consider during a split, as well as the minimum number of samples required to split and internal node or construct a leaf node. For Morgan fingerprints (cFP and bFP)^58^, we include an additional hyper-parameter to select for the optimal dimension of the fingerprints. In the end, we report the performance index using the best hyper-parameter choices from the scan, for each featurization.

### Performance index for mouse piriform cortical activity dataset

Following the original analysis for the dataset in Pashkovski et al^30^, we used correlation distance as the basis for measuring both neural activity distances and molecular representation (e.g. POM or Dragon^59^) distances for each molecule pair, after centering the values per-neuron or per-feature dimension. Pearson correlations were then used to measure how well each representation captured the neural activity distances observed in different experimental conditions (representing various parts of the brain and different sets of probes). The performance index is then calculated by averaging and rescaling these latter correlations across each experimental condition.

### Datasets used in Figure 1

Each of the datasets below is indicated with a letter ([x]) corresponding to its position in Figure 1b.

### Dataset for human olfaction

#### Dravnieks [c]

This dataset^28^ contains 128 unique molecules with 146 odor descriptor targets, where each molecule has a perceptual rating for each odor descriptor, which we can use for regression labels.

#### DREAM Olfaction Prediction Challenge (Keller et al.) [h]

This dataset is generated from the data published with the crowd-sourced DREAM Olfaction Prediction Challenge^12^ where we used the “gold” dilution (i.e. the one used to score the challenge) to generate the average perceptual rating for 369 molecules over 21 odor descriptor targets such as pleasantness, grass, garlic, sweaty, etc.

#### Human olfactory receptors [i]

This dataset is compiled from the literature and from databases as part of the OdoriFy effort^29^, and we use the binary response label for all eight different receptor targets including OR1A1, OR1A2, OR1G1, OR2J2, OR2W1, OR51E1, OR51E2, and OR52D1.

### Dataset for mouse piriform cortex activity [f]

The activity for each odorant and neuron pair is computed from the original raw time-series response curve kindly provided by the authors of a mouse piriform cortical activity dataset^30^. For each trial, the response curve is first smoothed by averaging each frame with a moving window of size 5. The baseline mean μ and standard deviation σ are then established using activities from the last 30 frames immediately before the designated odor onset. A response is elicited for the trial if the max response value within 30 frames after the onset is larger than μ + 3 * σ. The activity for each odorant and neuron pair is represented as the average elicitation rate across multiple trials.

### Dataset for insect olfaction

A total of 11 insect olfaction datasets are organized from 7 prior works in the literature and one previously unpublished data source.

#### MacWilliam et al.^22^ [a]

The authors perform a behavioral experiment with *Drosophila* using the T-maze assay. They measure the attraction and aversion with a preference index between −1 and 1 for around 60 compounds, as shown in Figure 5 of their paper. The wild-type preference index for each compound is extracted, and the dataset is represented as a binary classification task, where a compound is considered positive if it can elicit a strong attraction (>0.25) or a strong aversion (<−0.5).

#### Xu et al.^25^ [b]

The authors measure the odorant-elicited *in vitro* electrophysiological current response for CquiOR136 in the southern house mosquito, *Culex quinquefasciatus*. The authors measured this response for around 200 compounds as listed in experimental procedures. The compounds that can elicit detectable currents are found in Fig. 3 of that paper and are therefore assigned a positive label in this binary prediction task.

#### Mosquito repellency of fragrance molecules (Wei et al., unpublished) [d]

As part of an unpublished data source, 38 molecules were selected from a fragrance catalog, and their repellency was tested with a mosquito feeder assay: the fragrance molecules are coated on the membrane of a nano feeder containing 100 ul of blood meal. After feeding for 10 minutes, the average percentage of (%) unfed mosquitoes is measured over two trials. The repellency is then calculated by normalizing the unfed percentage such that ethanol has a 0% repellency, while the best possible repellency is 100%. This dataset is formulated as a binary classification task, where a compound is assigned a positive label if more than 90% repellency is observed.

#### Missbach et al.^23^ [e]

The authors perform single sensillum recordings for various types of olfactory sensory neurons (OSNs) in four different species including wingless bristletail, *Lepismachilis y-signata*, firebrat, *Thermobia domestica*, neopteran leaf insect, *Phyllium siccifolium*, and fruit fly, *Drosophila melanogaster*. The average spike count per second is recorded for a panel of 35 odorant molecules with six different functional groups. The spike count data is compiled for each species from Figure 3 of the paper, and formulated as a classification task where a compound is labeled as positive for a specific OSN target if the average elicited firing rate is 50% higher than the baseline firing rate.

#### del Mármol et al.^31^ [g]

The authors measure the odorant-elicited response for olfactory receptors MhOR1 and MhOR5 from the jumping bristletail, *Machilis hrabei*. The authors express the respective receptors in HEK cells and perform whole-cell recordings. An activity index is then computed for a panel of odorants based on the log(EC_50_) and the maximal response. The data from Supplementary Table 4 and 6 of that paper is compiled and formulated as a regression task with multiple targets.

#### Carey et al.^26^ [i]

The authors express various olfactory receptors for malaria mosquito *Anopheles gambiae* in ‘empty neurons’, and measure the olfactory receptor neurons (ORNs) responses as a consequence of odorant stimuli. We cast various receptor ORN responses as different regression targets, and compile the data from Supplementary Table 2c of the paper.

#### Hallem and Carlson^24^ [k]

The authors perform a similar assay as **Carey et al.**^26^, but with olfactory receptors in *Drosophila*. The data is extracted from Table S2 of the paper, and similarly compiled as a regression task with multiple targets.

#### Oliferenko et al.^27^ [n]

The authors measure the *Aedes aegypti* repellency for about 90 molecules as the minimum effective dosage (MED). Following the same threshold as the original paper, this dataset is casted as a binary classification task where compounds with an observed MED of less than 0.15 μmol/cm^2^ are considered active.

### Datasets for non olfaction related tasks

#### Enteric GPCR binding [q]

Five large human GPCR targets are pulled from GPCRdb^40^ including 5GT1A (serotonin receptor 1a), CNR2 (cannabinoid receptor 2), DRD2 (dopamine receptor 2), GHSR (ghrelin receptor), and OPRK (opioid receptor kappa). Since GPCRdb contains the binding affinities collated from multiple sources, we use the average binding score as our regression label for each target.

#### Other molecular properties [l, m, o, q, r]

From MoleculeNet^34^, five diverse tasks are selected as non olfaction related molecular properties, including electronic properties (QM7^39,60^), binding affinity with BACE-1 protein (BACE^38^), water solubility (ESOL^37^), hydration free energy (FreeSolv^36^), and adverse drug reaction (SIDER^35^). All these tasks contain a single regression label except SIDER which is a multi-label classification task.

### Dataset Standardization

All olfactory datasets are standardized by removing the following molecules: 1) data points with multiple molecules (i.e. mixtures), 2) molecules with only a single atom, 3) molecules with atoms that are not hydrogen, carbon, nitrogen, oxygen and sulfur, or 4) molecules with a molecular weight of more than 500 daltons. Nearly all odorant molecules in the raw datasets passed this empirically-motivated standardization filter^43^ and are kept in the standardized dataset.

### Metabolic networks and metabolic distance

A metabolic network is constructed for each species where each node represents a metabolite and each directed edge connects the reactant and product metabolites in different experimentally elucidated reactions in the MetaCyc database^2^. Among all metabolic networks for different species, the 17 largest networks each with more than or equal to 100 metabolites are further studied.

For each metabolic network, all metabolites are labeled odorous or not according to mass transport principles established in Mayhew et al., 2022^43^. All pairs of odorous metabolites with an existing path in their network are enumerated and the distance of their shortest path is used as their metabolic distance — the minimum number of metabolic reactions to convert one odorous metabolite to another. Due to the sparsity of these metabolic networks, far more metabolite pairs with short metabolic distances (<3) are found compared to those with long metabolic distances (>8). In order to fairly study the organization of metabolite pairs across various distances, metabolite pairs are resampled such that an even number (=50) of metabolite pairs are sampled for each metabolic distance. The Pearson correlation coefficients shown in Figure 2 are also consistent across different sampling seeds (Extended Data Figure 5) and are destroyed by perturbations that corrupt the metabolic graph (Extended Data Figure 6).

### Computing distances

The POM distance between molecules is defined as the correlation distance between their embeddings, which are centered across the population of molecules (e.g. all sampled odorous metabolite pairs, or all compounds in essential oil). Tanimoto distance^33^ is computed using RDKit^61^ based on bit-based fingerprints. cFP edit distance between two molecules approximates the absolute difference between their structures and is defined as the L1 distance between their count-based fingerprints.

### Visualizing metabolic pathways and calculating their smoothness

In order to visualize the metabolic pathways (Figure 3a,b), both the count-based fingerprints (cFP) and principal odor map (POM) are projected to a compressed 64 dimensional space via principal components analysis. Using subspaces with a common number of dimensions also controls for any dimensionality bias that might be present in these comparisons. Explaining more than 80% of the variance in both representations, the first two dimensions of the PCA projections are then used to visualize the trajectory of these pathways in cFP vs. POM.

To quantify the “smoothness” of these metabolic pathways (Figure 3d), 34 unique metabolic pathways with distance ranging from 3 to 13 are extracted from the 15 metabolic networks. For an example pathway “A->B->C->D”, we can calculate the smoothness for interpolation of the two intermediate metabolites B and C. The smoothness for interpolating B can be defined as **d**(A, D) / (**d**(A, B) + **d**(B, D)), where **d** is the euclidean distance between the 64 dimensional PCA projection of cFP or POM (Figure 3c). In total, 61 such valid odorous metabolite triplets are found and evaluated.

### Common scents for molecule pairs

The common scents for molecule pairs listed in Figure 2d and Figure 4d are predicted by a state of the art model^41^. An odor label is assigned to the molecule if the model predicts a label probability > 0.5 for that label. Multiple labels are possible for each molecule.

