## Supplemental Materials for "Metabolic activity organizes olfactory representations"

Extended Data Figures (Supplementary Figures)

***Extended Data Figure 1. Performance index for alternative structure-based featurizations.*** **a)** Similar to Figure 1b, but showing performance of both POM (gray) and alternatives (count-based fingerprints, cFP, red; and bit-based fingerprints, bFP, orange) against Mordred chemoinformatic features. Only POM shows a consistent advantage in olfactory tasks. **b)** Summary of a) by averaging the performance ranking of different featurization methods, grouped by the origin of the task. Here, a higher rank means better performance — the best rank is 4 and the worst rank is 1.

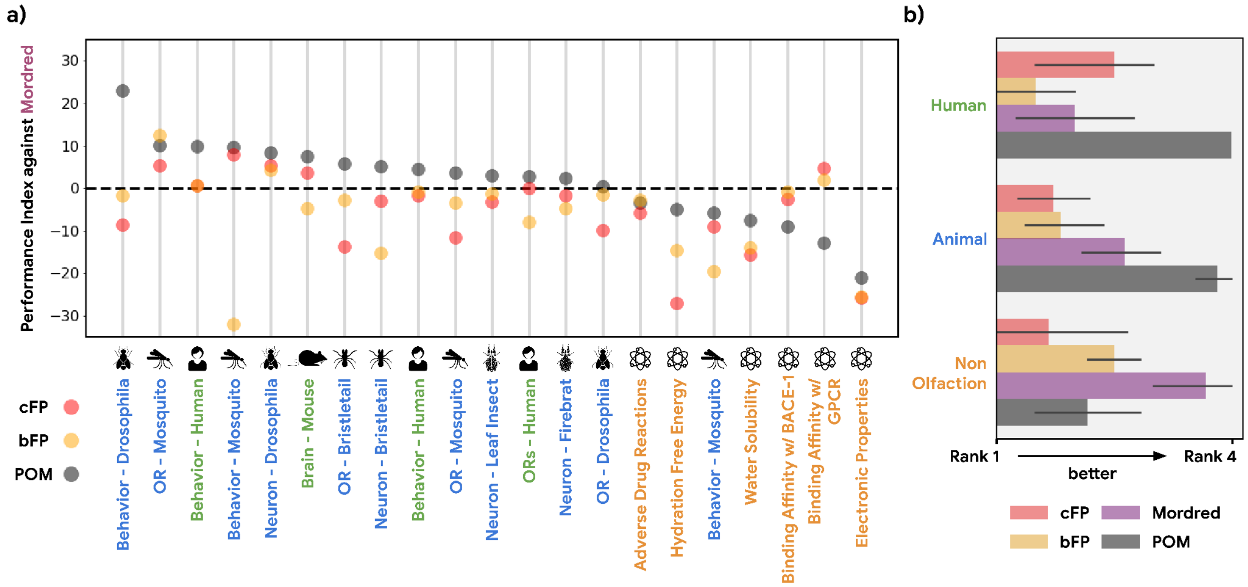

***Extended Data Figure 2. Relative performance of POM is a function of dataset distance.*** The advantage in predictive performance for POM vs. Mordred (i.e., see vertical axis in Fig. 1b) decreases as the dataset becomes less relevant to olfaction ($r=0.49$, $p<0.03$). Relevance is measured by dataset distance (farther is less relevant), quantified as the distance from the training data (human odor labels) computed via the optimal transport method [1].

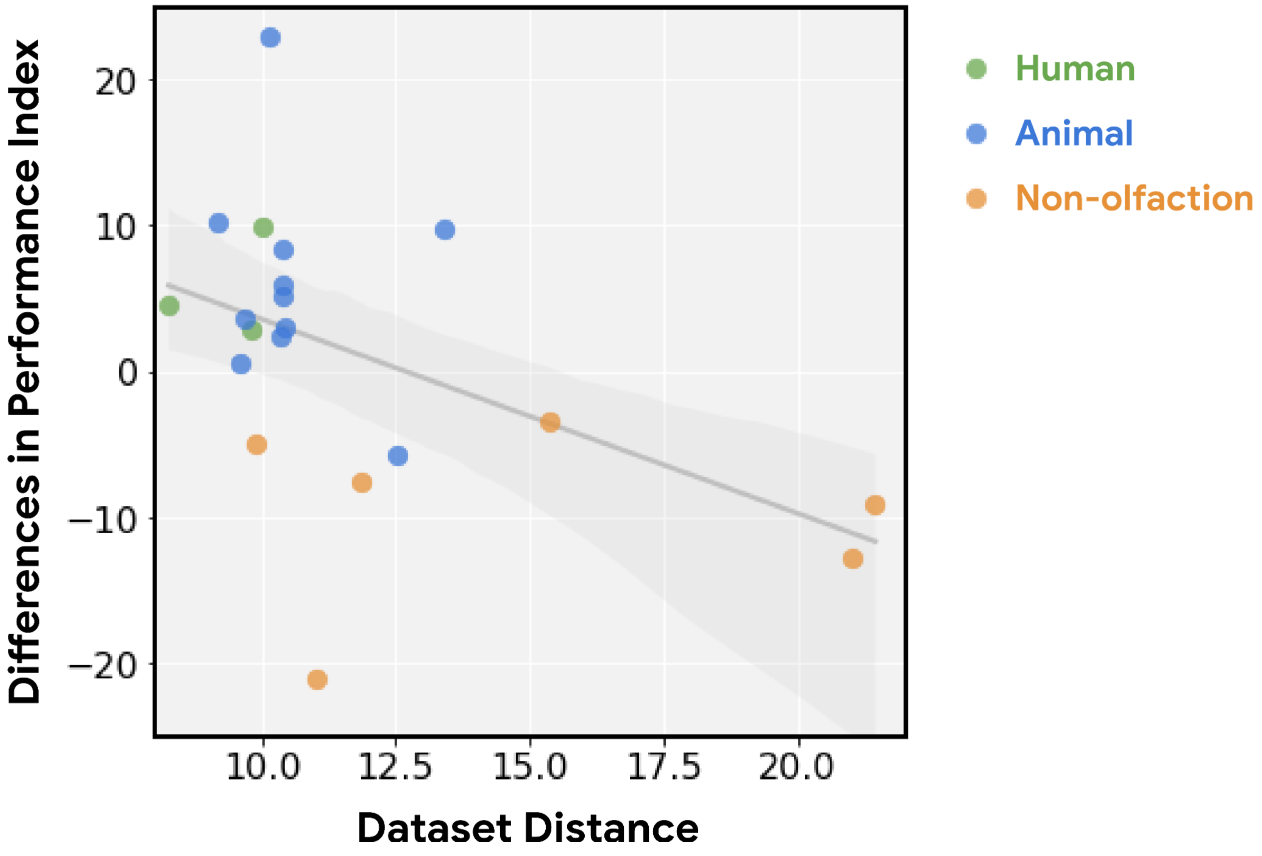

***Extended Data Figure 3. POM has no predictive advantage for non-olfaction-related chemoinformatic properties of odorous molecules.*** In Figure 1 and Figure ED1, we estimate the performance index of POM and other methods on non-olfaction datasets containing molecules that are not necessarily odor-molecule-like. In order to control for such molecular distribution shifts, we construct four datasets for non-olfaction-related tasks on the same set of odorous molecules used in the training dataset. For general molecular properties, we compute Wildman-Crippen LogP[2], ESOL[3], and QED[4] values for molecules in the training dataset; for enteric GPCR binding, we estimated the binding affinity between these molecules and the same GPCR used in Figure 1 using the winning model from the IDG-DREAM Drug-Kinase Binding Prediction Challenge[5]. As in Figure 1, POM shows no advantage on these non-olfaction tasks, and is strictly dominated by Mordred across each such task.

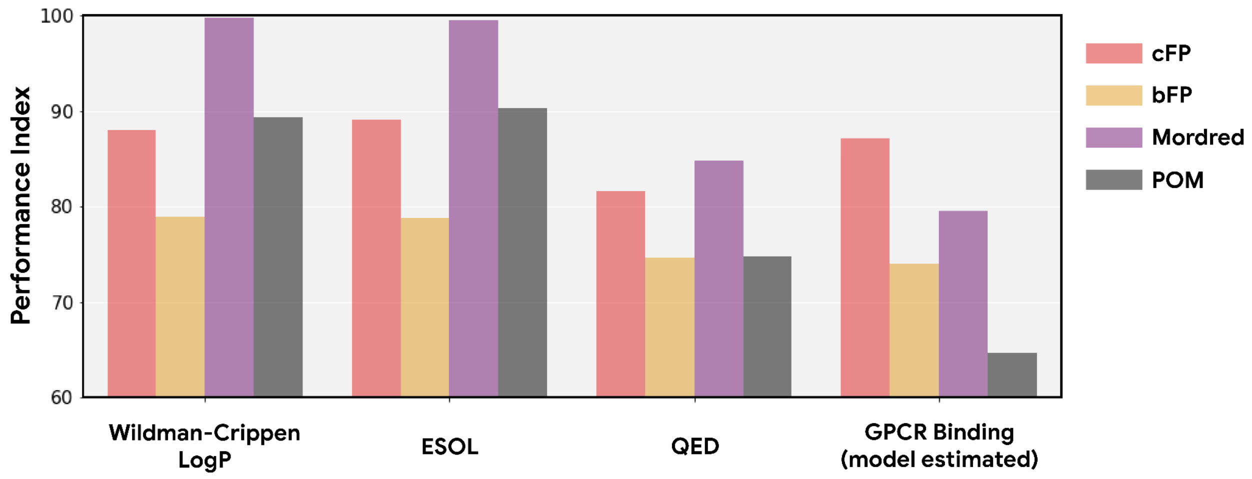

***Extended Data Figure 4*. Distributional statistics for metabolic networks. a)** Distribution of the count of metabolites described in MetaCyc across individual species. Most species were not well-represented, resulting in sparse networks, but 17 species were represented by >=100 metabolites which we then selected for the analysis. **b)** Most metabolites did not pass a simple odorousness filter, but we still find 405 pairs of odorous metabolites that can be connected via various steps of metabolic reactions. **c)** Most odorous metabolites were connected by a short reaction path (<=3 steps), but the path can contain as many as 12 metabolic reactions. **d)** The 17 species whose metabolic networks are used here span four different kingdoms of life.

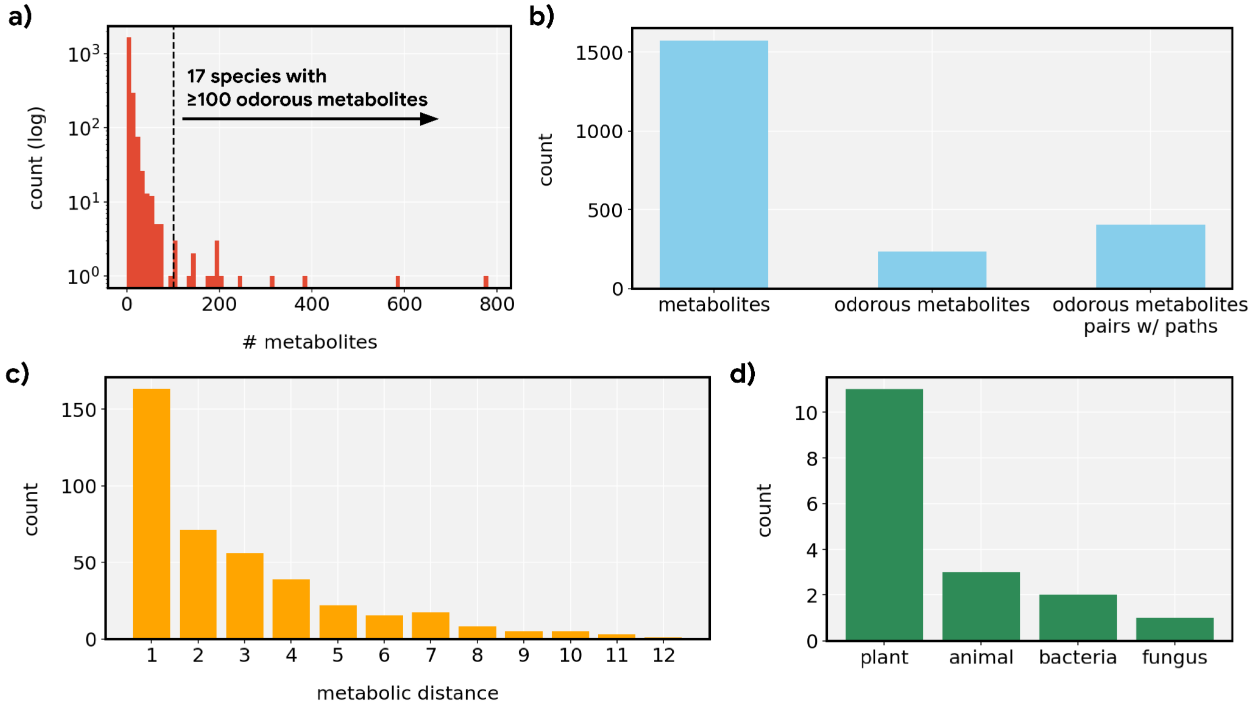

***Extended Data Figure 5*. Correlations are robust to alternative sub-samples of metabolite pairs.** Box plots show the median, inter-quartile range, and min/max correlations for 100 sampling seeds, i.e., for alternative sub-samples of metabolite pairs compared to those shown in Figure 2c. The correlation for POM distance is uniformly stronger than for other distance measures.

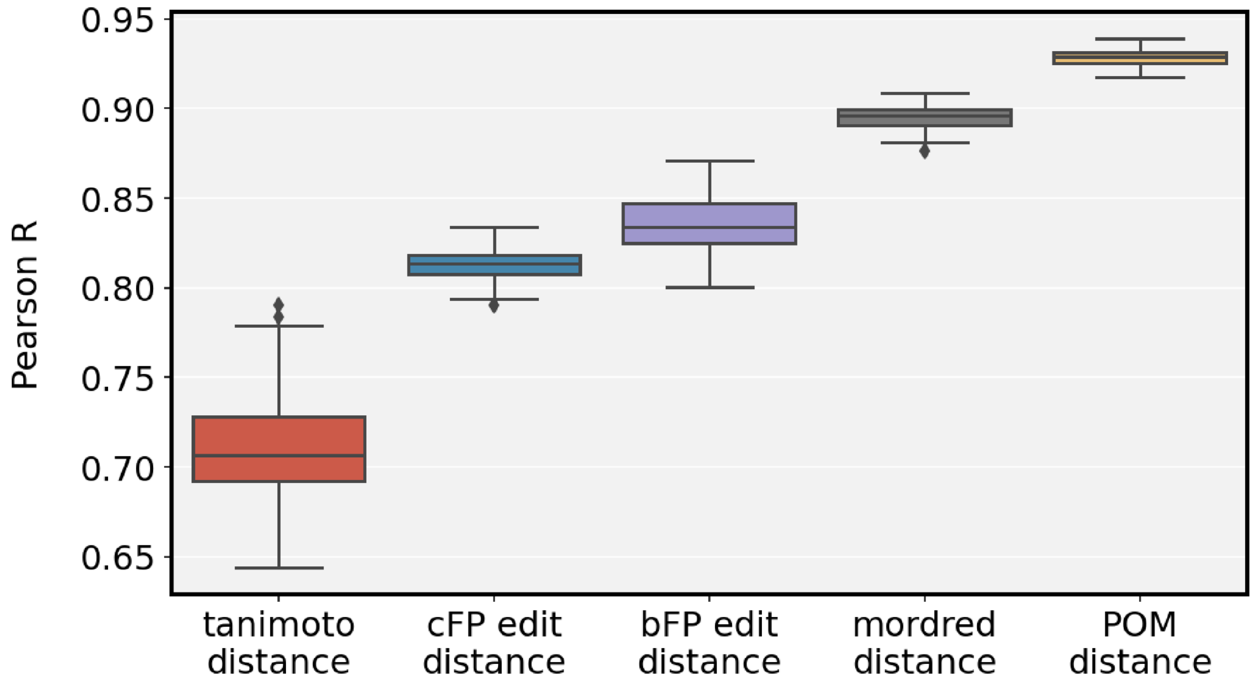

***Extended Data Figure 6*. Perturbation of the metabolic graph destroys the observed relationships.** Correlations (across random seeds as shown in Figure S5) between metabolic distance and the indicated intermolecular distance, for four different perturbations on the original metabolic networks: **a)** End points of all the edges are randomly re-assigned to either the original metabolite or its neighbors**; b)** Directions of 50% of the edges are reversed (i.e. reactants become products and vice versa); **c)** Ablation of direction information (reactant vs product identity becomes indeterminate); **d)** Merging of all graphs across organisms, i.e. any reaction in an organism can be used to determine metabolic distance.

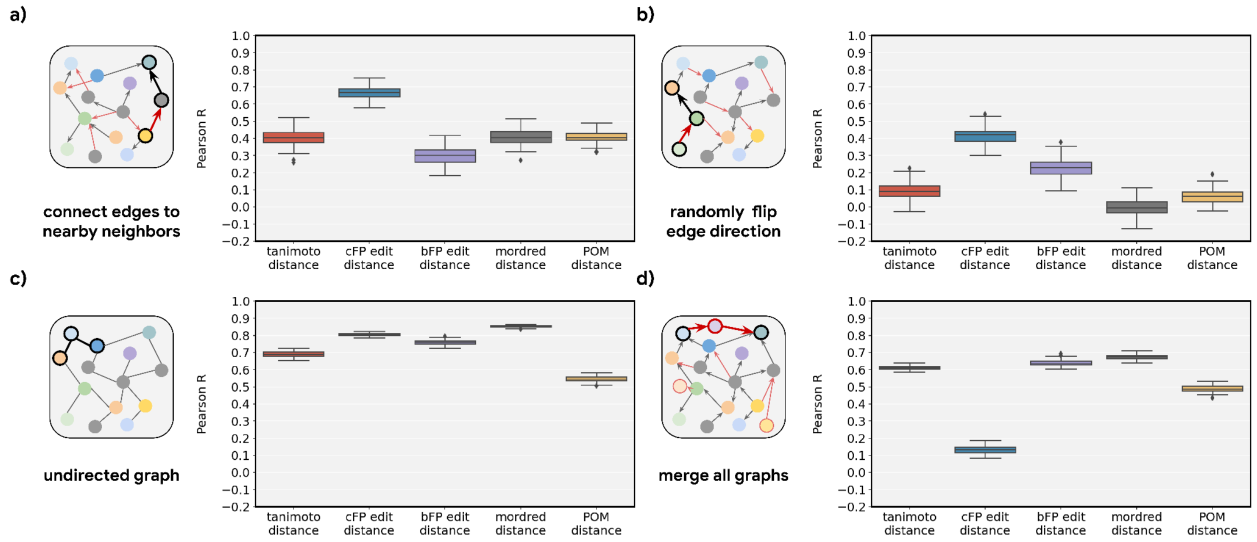

***Extended Data Figure 7*. The smoothness of metabolic pathways in POM is not an artifact of dimensionality. a)** Cumulative explained variance as a function of the number of principal components for different featurizations. The inherent dimensionality of POM is lower than the alternatives. **b)** The smoothness metric reported in Figure 3d, as a function of the number of principal components retained. The dashed line (64 components), reflects the value used in Figure 3d. Reducing the dimensionality to a common value was necessary to avoid over-penalizing higher-dimensional feature spaces (e.g., bFP, cFP, Mordred).

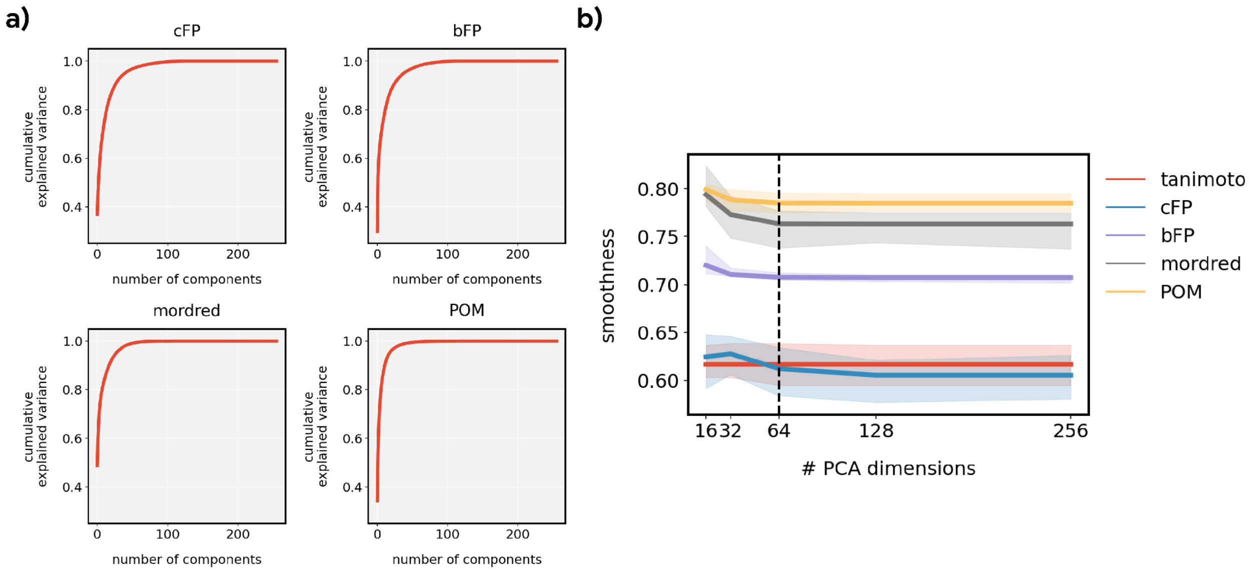

***Extended Data Figure 8.* Smoothness of individual metabolite triplets.** The metabolite triplets analyzed in Figure 3d are shown here in detail; each circle represents the smoothness of one triplet (e.g., A->B->D or A->C->D). Most triplets are smoother for POM than for alternative representations (i.e., circles are above the dashed line). The smoothness across all triplets is significantly higher for POM than for each of the alternatives (paired t-test; p-value shown in each panel).

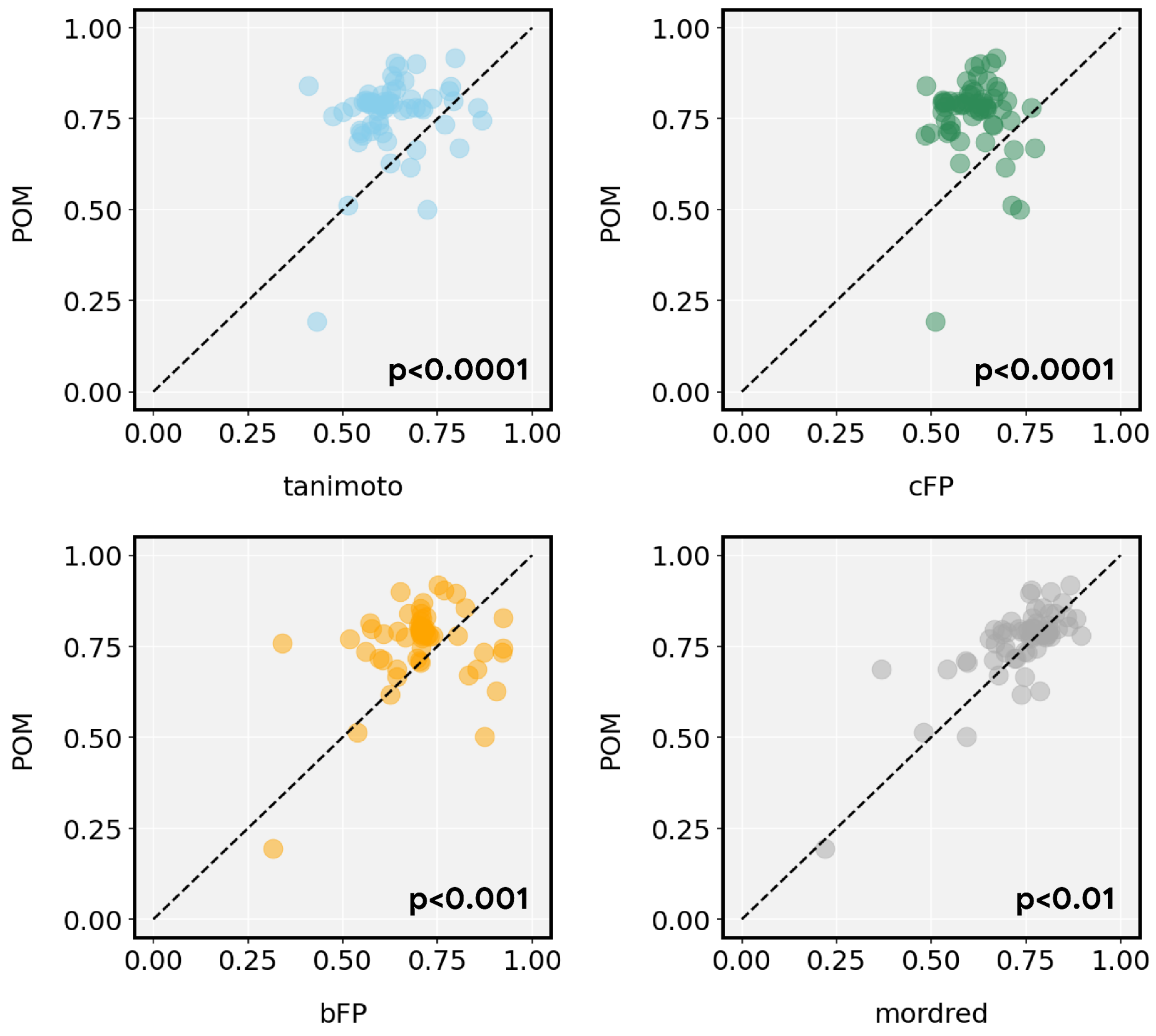
